# Allelic diversity study of functional genes in East Africa bread wheat highlights opportunities for genetic improvement

**DOI:** 10.1101/2020.05.11.088104

**Authors:** Mercy Wamalwa, Zerihun Tadesse, Lucy Muthui, Nasser Yao, Habtemariam Zegeye, Mandeep Randhawa, Ruth Wanyera, Cristobal Uauy, Oluwaseyi Shorinola

## Abstract

Wheat (*Triticum aestivum* L.) is a major staple crop in East Africa (EA) providing 9% and 10% of daily calories and protein intake, respectively. However, EA countries depend on import to meet 55% of their domestic wheat supplies due to increasing demands and low domestic yields. To determine the beneficial gene pool currently exploited for wheat improvement in EA, we examined the allelic diversity of 42 genes of breeding importance in a collection of 239 wheat cultivars and breeding lines from Kenya and Ethiopia using KASP markers. The assayed genes have been shown to control variations in plant height, thousand kernel weight (TKW), grain protein content, pre-harvest sprouting (PHS), disease resistance and flowering time. We observed the beneficial alleles of some major genes including *Rht-D1, Gpc-B1, Yr5, Yr15, Sr26*, and *Fhb1* to be missing or present at low frequencies in this population. Furthermore, we validated the effects of the major *Rht-1* alleles and *TaCKX-6A* in controlling variation in plant height and thousand kernel weight, respectively, under EA conditions. Our results uncover hitherto unexploited allelic diversity that can be used to improve the genetic potential of EA wheat germplasm. This will inform strategies to rapidly mobilize these beneficial alleles for wheat improvement in EA.

## Introduction

Wheat (*Triticum aestivum* L.) is one of the major cereals and staple crop in East Africa (EA). Due to the increasing rates of population growth and urbanization, as well as increases in household incomes in many EA countries, there has been a shift in dietary pattern to easy-to-cook meals including wheat-based diets (Negassa et al. 2013). This has led to a two-fold increase in per-capita wheat consumption in EA over the last 60 years, at a time when percapita consumption is declining in Europe. Wheat has therefore transitioned from a “colonial” crop confined to missionaries shambas (farms) to a food security crop accounting for 9% and 10% of daily calorie and protein intake in EA, respectively (FAOSTAT 2020). Up to 80% of wheat farmers in EA are small holder farmers, further underlying the importance of wheat production to household food security and livelihoods. Kenya and Ethiopia account for 92% of the wheat produced in EA, however, the average yield of wheat in both countries is low (2 tonne/ha in 2017; FAOSTAT 2020). To meet the increasing demand for wheat, EA countries import 55% of their domestic wheat supplies. This heavy dependence on imports makes the region highly vulnerable to changes in global wheat supply market.

Advances in wheat genomics have enabled the identification of genes, and their allelic variations, that control many economically important traits in wheat. The development of molecular breeding approaches has also paved way for deploying beneficial alleles of these genes to accelerate wheat improvement in national and global wheat breeding programs. There is a long history of wheat breeding in Kenya and Ethiopia, however, the use of molecular breeding tools is very limited thereby hampering the rate of genetic gains achieved. As such, the national breeding programs in both countries have depended on introductions of wheat lines from international wheat breeding programs including CIMMYT and ICARDA. Understanding the composition and diversity of the gene pool in the EA wheat germplasm is important for defining breeding strategies and prioritizing trait targets for wheat improvement.

Kompetitive Allele-Specific PCR (KASP) is regarded as a benchmark for SNP genotyping in marker assisted breeding due to its low-cost, ease-of-use, specificity and co-dominant nature (Semagn et al. 2014). Over the last decade, many functional KASP markers have been developed for genotyping allelic variations controlling economically important traits in global wheat germplasm. Functional genetic markers, unlike random genetic markers, are derived from allelic variations that have been shown to affect phenotypic variation either by association, linkage or causation (Andersen and Lübberstedt 2003). There have been concerted efforts to assemble functional KASP markers in databases and publications making them easily accessible for marker-assisted breeding (CerealsDb: Wilkinson et al. 2012; MASWheat: https://maswheat.ucdavis.edu/). Recently, Rasheed et al. (2016) and Khalid et al, (2019) together describe the assembly of 127 functional KASP markers for studying allelic variation in 88 genes underpinning agronomic, adaptation, end-use quality, disease resistance and drought tolerance traits.

The aims of this study were to: (1) use readily-available KASP markers to evaluate the allelic diversity of genes of breeding importance; (2) examine the changes in the allele frequencies of these genes over the last century; (3) evaluate the phenotypic effect of some of these genes on plant height and grain size variations under EA field conditions.

## Materials and Methods

### Plant materials and phenotyping

A set of 239 bread wheat genotypes comprised of 150 lines from Ethiopia and 89 lines from Kenya were used. The Ethiopian wheat genotypes were obtained from the Ethiopian Institute of Agricultural Research (EIAR) and comprised 53 wheat cultivars released from 1970 - 2017 and 97 breeding lines currently under development. The Kenyan wheat genotypes comprised cultivars released from 1920 to 2012 and were obtained from the Kenya Agricultural and Livestock Research Organization, (KALRO).

Three phenotyping experiments (Experiment 1 - 3) were conducted in Kenya and Ethiopia across three years. Experiment 1 and 2 were conducted independently for Kenyan and Ethiopian genotypes, respectively, to account for local environmental effects on the performance of genotypes from each country. Experiment 1 was conducted at the EIAR Kulumsa Agricultural Research Centre (8° 01’N, 39°09’ E; 2,200 m above sea level) during the 2017 main cropping season from July to November. Each line was sown in double row plots (1 m) using alpha lattice design with two replications. Experiment 2 was conducted at KALRO Field Crop Research Station Njoro (0° 20’S, 35° 56’E; 2,185 m above sea level) during the 2018 main cropping seasons from May to October. Experiment 3 comprised both Kenya and Ethiopia genotypes and was conducted at the KALRO Field Crop Research Station in 2019. Each entry was sown in double rows measuring 0.2 × 0. 75 m.

Plant height measurements were obtained across all three experiments. Plant height was measured at physiological maturity from the plant base to the tip of the spike (excluding awns) of the primary tiller. For Experiment 1 and 2, each plot was hand-harvested at maturity, threshed, and a sample of 1000 kernels was taken and weighed to determine the Thousand Kernel Weight (TKW).

The year of release and phenotypic data for each genotype are presented in Table S1 in Online Resource 1.

### DNA extraction and KASP genotyping

Leaf samples were collected from single plant for each line at the seedling stage. Genomic DNA was extracted following the CTAB method (Doyle and Doyle 1987) and ZYMO kits (ZR Plant/Seed DNA mini-Prep kit; Zymo Research, USA). The quality and quantity of DNA extracted was evaluated using spectrophotometric (NanoDrop^™^ 8000) and gel-based methods. All samples were diluted to a uniform concentration, and an equivalent of 50 ng of DNA per sample was used for genotyping.

KASP genotyping was done at the BecA-ILRI Hub (Kenya). A total of 59 KASP assays were used in this study. This comprises 55 of the 127 assays reported by Rasheed et al (2016) and Khalid et al (2019). We also obtained assay information for *TaMKK3-A, Yr15, Yr5* and *Sr26* directly from the publications describing the development of KASP markers for these genes (Shorinola et al. 2017; Klymiuk et al. 2019; Marchal et al. 2018; Qureshi et al. 2018). Each assay comprises of two allele-specific primers tailed with either the FAM or HEX dye sequences and a common untailed primer. The gene information, primer sequences and allele-dye assignment for each marker used are presented in Table S2 in Online Resource 1. The primers were mixed in the following proportion: 46 μL de-ionized water, 30 μL common primer (100 μM), and 12 μL of each tailed primer (100 μM). KASP genotyping was performed following the manufacturer (LGC Biosearch Technologies, UK) protocol. The assays were run in 384-well plate format. A reaction volume of 5.0 μL comprising 2.5 μL of DNA and 2.5 μL of a mix of 2x KASP reagent and 0.07 μL primer mix was used for amplification. PCR cycling was performed using the following protocol: hot start at 94 °C for 15 min, followed by ten touchdown cycles (94 °C for 20 s; touchdown at 61 °C initially and decreasing by −1 °C per cycle for 60 s), followed by 30 cycles of annealing (94 °C for 60 s; 55 °C for 60 s). Additional cycles were performed to increase the intensity of the fluorescence signals where necessary. For each assay, we included no-template (water) controls as well as varietal controls that harbor alternate alleles of each gene (where available). Fluorescence signals from amplified samples were measured at <40 °C using the Fluostar Omega Microplate Reader (BMG Labtech, UK). Genotype clustering was done using the KlusterCaller software (LGC Biosearch Technologies, UK).

### Statistical Analyses

Statistical analyses were done in R 3.6.0 (R Core Team 2019) using the LmerTest and car packages (Kuznetsova et al. 2017; Fox and Weisberg 2019). Only allelic variations with minimum allele frequency > 0.05 were included for statistical analyses. As plant height measurement was taken in at least two experiments for each genotype across three years, a linear mixed-effect function using Restricted Maximum Likelihood model was used to assess the effect of *Rht-1* genes on plant height. For this, allelic variation of *Rht-1* genes, year (2017 – 2019), and location (Kenya or Ethiopia) were taken as fixed effects while genotype was taken as a random effect to account for the background genetic effects from each genotype. TKW measurement was only taken at one location (in Experiment 1 or 2) for each genotype. A linear model taking allelic variation for the grain size genes and location (Kenya or Ethiopia) as fixed effects was, therefore, used to assess the effect of grain size related genes on TKW.

## Results

### Allele frequencies of functional genes in EA wheat population

We genotyped 239 EA bread wheat cultivars and breeding lines using 59 KASP assays. Of these 59 assays, we obtained good genotype clusters (distinct from the no-template control) for 48 assays. These 48 assays target 42 genes controlling plant height, drought tolerance, grain quality, grain size, flowering time, and disease resistance. Genotype information of all 48 working assays are presented in Table S3 in Online Resource 1. The remaining assays either produced clusters that were difficult to separate or had high proportions of samples with no amplification. Of the 48 working assays, 14 assays produced only one distinct genotype cluster separate from the no-template control (Figure 1A and B) suggesting that the germplasm are fixed for a single allele. We considered a gene allele to be favorable if it is causal, associated or linked to a desired agronomic, disease resistance, grain size or end-use quality traits (Figure 2, see details in Table S2). Ten of the 14 single-cluster assays were fixed for unfavorable gene alleles, thus highlighting immediate opportunities for improving these traits in the EA wheat population. Some of the major genes fixed for unfavorable gene alleles in the EA wheat population include *Yr5, Fhb1* and *Sr26*.

**Figure 1:**
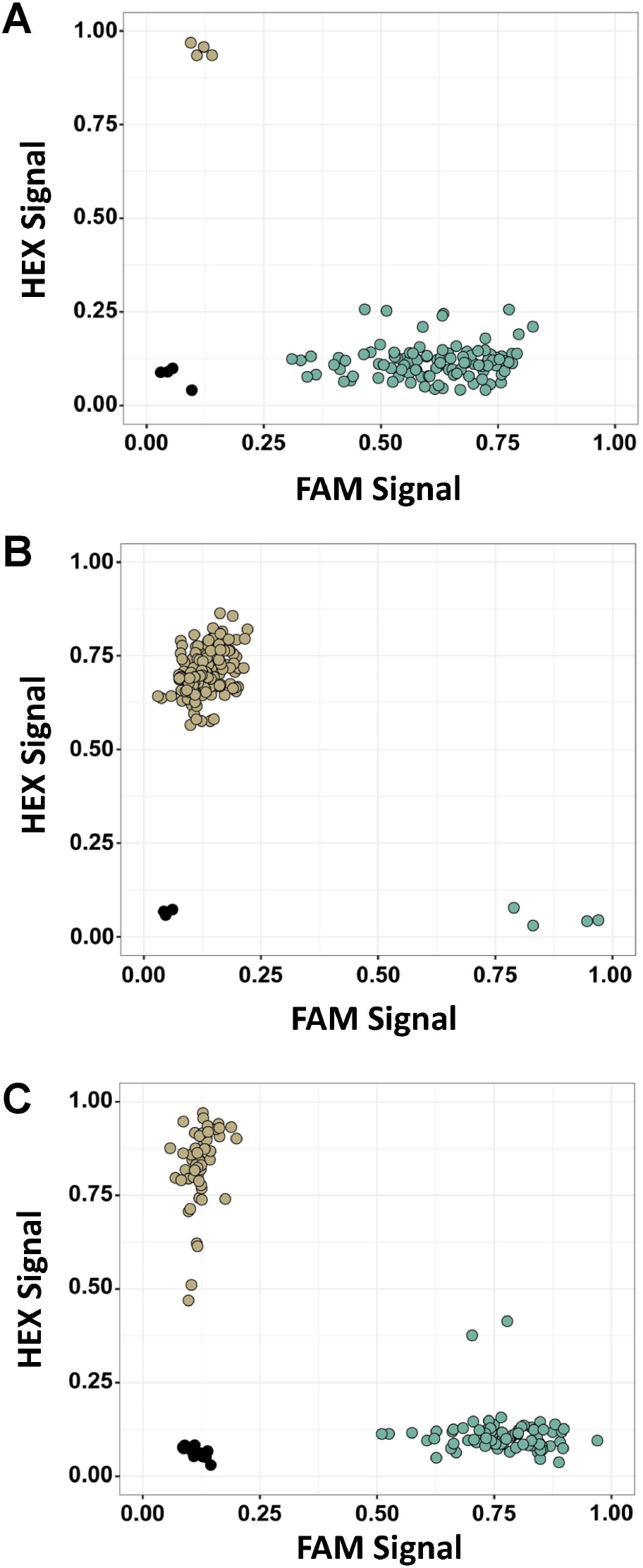
KASP marker clustering pattern. Representative genotype clustering pattern of functional KASP markers in EA wheat population showing fixation for a favorable allele (a), unfavorable allele (b) or segregation for both alleles (c).

**Figure 2:**
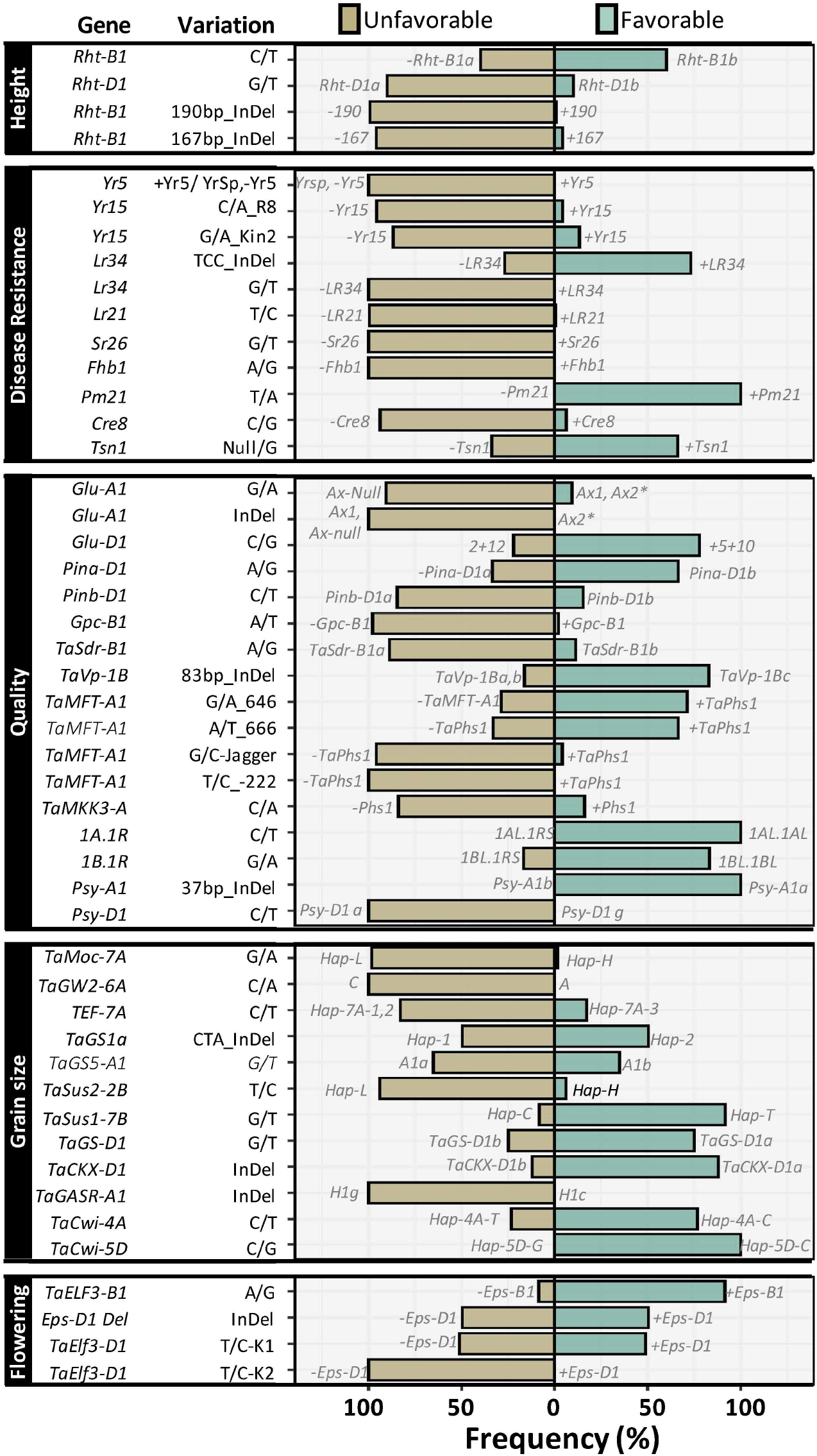
Allele frequencies of functional KASP markers in EA population. Allele frequencies of 48 markers for height, disease resistance, grain quality, grain size and flowering time genes in EA population. Polymorphism information for each gene is described on the left. The frequencies of favorable and unfavorable gene alleles are represented with green and lightbrown colored bars, respectively. The allele nomenclature for each gene are indicated next to each bar.

**Figure 3:**
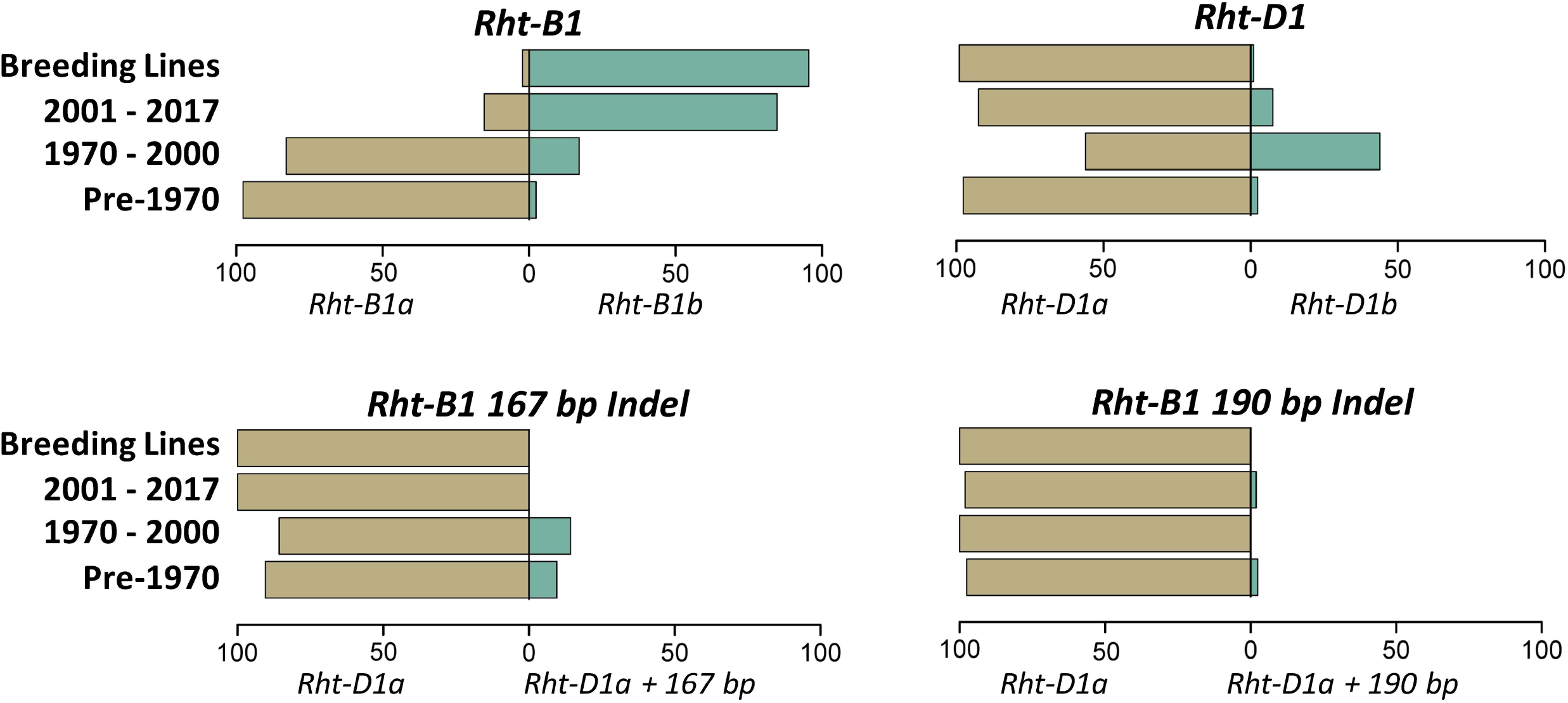
Frequency distribution of plant height gene alleles. The frequencies of plant height gene alleles across four breeding periods are represented. Frequencies of the favorable and unfavorable gene alleles are represented with green and light-brown colored bars, respectively.

**Figure 4:**
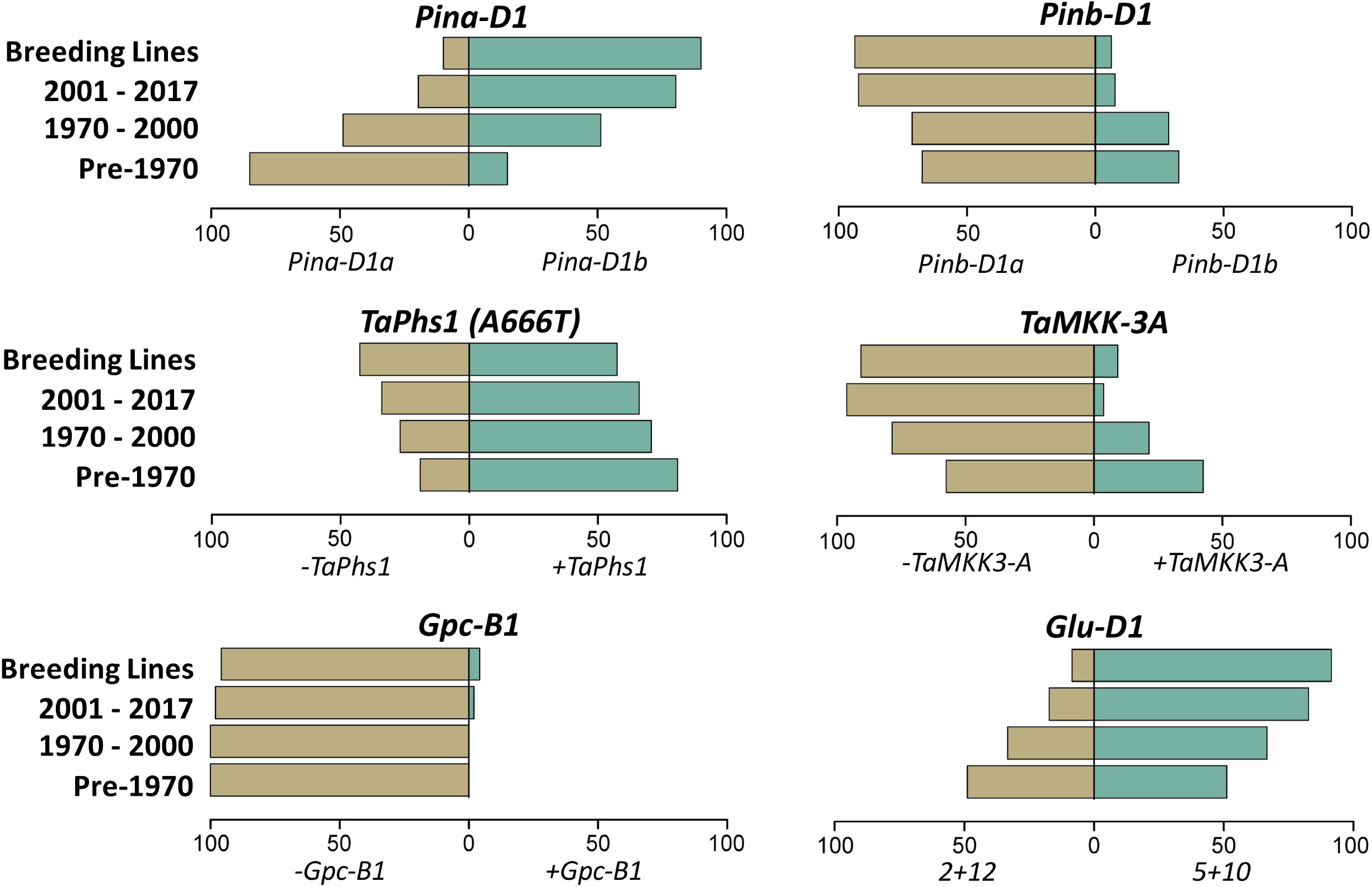
Frequency distribution of grain quality genes alleles. The frequencies of grain quality gene alleles across four breeding periods are represented. Frequencies of the favorable and unfavorable gene alleles are represented with green and light-brown colored bars, respectively.

**Figure 5:**
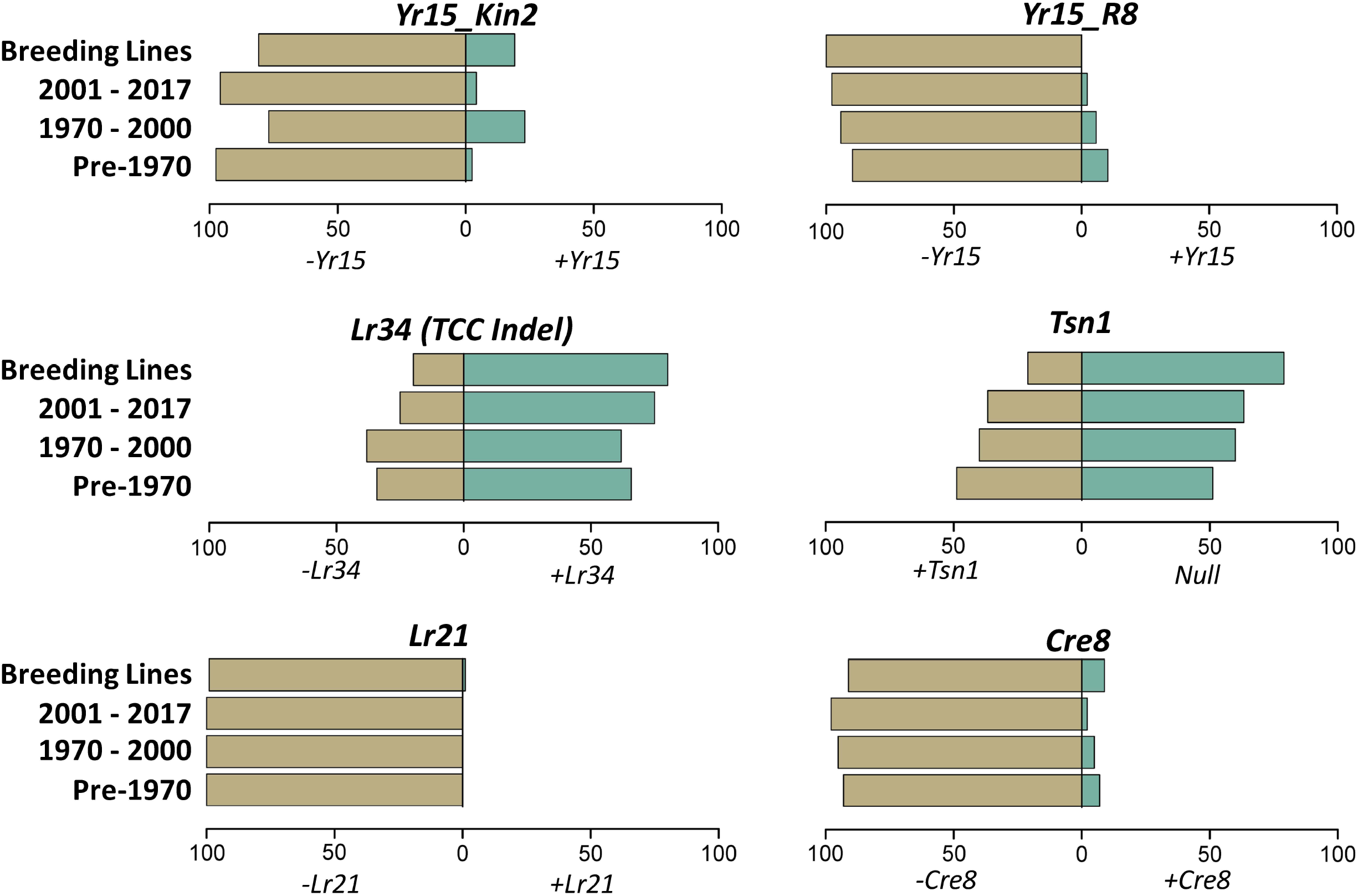
Frequency distribution of disease resistance gene alleles. The frequencies of the disease resistance gene alleles across four breeding periods are represented. Frequencies of favorable and unfavorable gene alleles are represented with green and light-brown colored bars, respectively.

On the other hand, 34 assays produced two distinct genotype clusters suggesting segregation of favorable and unfavorable alleles (Figure 1C). Of these segregating assays, 14 assays showed higher frequencies for the favorable alleles, while 17 showed higher frequencies for the unfavorable alleles (Figure 2). Three assays showed near-balanced frequencies between the favorable and unfavorable alleles.

Allele frequencies distribution for most of the genes examined showed a similar pattern in the Ethiopian and Kenya germplasm in terms of pre-dominance of the favorable or unfavorable alleles (Figure S1, Online Resource 2). This is most likely due to the common sources of founder materials and the frequent exchange of germplasm between the national wheat breeding programs of both countries. The only exceptions were for *Rht-B1* and *Pina-D1*. The favorable allele of *Rht-B1* was pre-dominant (86%) in Ethiopia compared to only 21% in Kenya. Similarly, the favorable allele of *Pina-D1* was pre-dominant (85%) in Ethiopia compared to only 37% in Kenya. The difference in allele frequencies for *Rht-B1* is most likely due to the composition of the wheat collection from both countries in our study: 50% of the Kenya collection were released before the introduction of *Rht-1* genes during the Green Revolution while the Ethiopian collection comprised entirely of post-Green Revolution lines.

### Changes in allele frequency distribution across breeding periods

The germplasm examined were introduced to or developed in EA across a 100-year period (1920 to 2017) spanning different breeding periods with access to different gene alleles and breeding technologies. As such we further examined the change in allele frequency distribution across these breeding periods for assays showing segregation. Four distinct wheat breeding periods were defined: pre-1970, before the introduction of the Green Revolution wheat cultivars to EA; 1970 – 2000, characterized by the introduction of Green Revolution derived germplasm into EA; 2001 – 2017, characterized by the use of molecular breeding tools; and breeding lines that are yet to be released. Due to their similar allele frequency distribution, the Kenya and Ethiopian germplasm released in these breeding periods were considered together. We focused mainly on the distribution of allelic variations that are either causal or linked to target phenotypes as these are most likely to have contributed to trait variations and breeder selection compared to associated allelic variants that are yet to be validated. The distribution of associated allelic variations (mainly for grain size genes) are presented in Figure S2 (Online Resource 2).

#### Plant height genes

Four assays for the major reduced plant height gene alleles, *Rht-B1* (*Rht-B1a:* tall; *Rht-B1b:* semi-dwarf), *Rht-D1* (*Rht-D1a:* tall; *Rht-D1b:* semi-dwarf) and the 167 bp and 190 bp InDel polymorphisms in *Rht-B1* were examined (Ellis et al. 2002; Wilhelm et al. 2013). *Rht-B1* was the pre-dominant gene conferring reduced plant height in the EA wheat population, with 60% of EA wheat lines harboring the favorable semi-dwarf *Rht-B1b* allele compared to only 10% of EA lines containing the semi-dwarf *Rht-D1b* allele. Furthermore, the allele frequencies of *Rht-B1b* increased in lines released post-1970 with a drastic increase in lines released from 2001. While the allele frequency of the favorable semi-dwarf *Rht-D1b* initially increased in lines released post-Green Revolution, the frequencies subsequently decreased sharply in lines released since 2001. None of the genotypes examined harbor *Rht-D1b* and *Rht-B1b* alleles together. The 167 bp and 190 bp polymorphisms show no substantial change in allele frequencies distribution across the breeding periods.

#### Grain quality genes

Six grain quality genes showed progressive changes in allele frequency distributions across the breeding periods. These included the puroindolines (*Pins*) genes, *Pina-D1* and *Pinb-D1*, which together control grain hardness (Morris 2002). Due to the higher premium for hard wheat grains compared to soft wheat grains, alleles that promote grain-hardness (*Pina-D1b* and *Pinb-D1b*) were considered favorable relative to alleles for grain-softness (*Pina-D1a* and *Pinb-D1a*). Grain-hardness is determined by the presence of either or both *Pina-D1b* and *Pinb-D1b*. We observed a steady increase of the favorable *Pina-D1b* allele across the breeding periods from 15% in lines released before 1970 to 90% in current breeding lines. Conversely, the frequency of the favorable *Pinb-D1b* allele decreased across the breeding periods from 33% in lines released before 1970 to 7% in breeding lines. We also examined the distribution of the *Pina-D1/Pinb-D1* haplotype across the breeding periods. The soft-grain *Pina-D1a/Pinb-D1a* haplotype decreased steadily across the breeding periods as did one of the hard-grain haplotypes (*Pina-D1a/Pinb-D1b*). In contrast, the *Pina-D1b/Pinb-D1a* hard-grain haplotype increased across the breeding periods and is the pre-dominant haplotype in the breeding lines (Figure S3, Online Resource 2).

*TaMFT-A1* (same as *TaPhs-A1*) and *TaMKK3-A* are two important genes controlling embryo-imposed seed dormancy in wheat (Vetch et al. 2019). Optimized seed dormancy level is advantageous for preventing pre-harvest sprouting (PHS) without compromising uniform seed germination. Different allelic variation exists for *TaMFT-A1* including a promoter polymorphism (−222 bp C/T; Nakamura et al. 2011) which is fixed in the EA population, and two segregating exonic and splice junction polymorphisms (A666T and G646A; Liu et al. 2013). Only one causal natural allelic variation has been reported for *TaMKK3-A* (Torada et al. 2016). We found higher frequencies of dormancy-promoting alleles of *TaMFT-A1* compared to *TaMKK3-A* in the EA wheat population. These frequencies of the dormancy-promoting *TaMFT-A1* alleles however reduced over the breeding periods. The frequencies of the dormancy-promoting *TaMKK-3A* allele showed a similar progressive decrease across the breeding periods.

Other segregating grain-quality genes examined included *Gpc-B1* (grain protein content) and *Glu-D1* (gluten strength). The high-protein *Gpc-B1* introgression was missing in varieties released before 2000. It is, however, present at low frequencies in varieties released post-2000 (2%) and in breeding lines (4%). We also observed an increasing trend for the strong-gluten *Glu-D1* (*Dx5 + Dy10*) alleles from 51% in pre-1970 lines to 92% in breeding lines.

#### Disease resistance genes

Only six of the 11 disease-related allelic variations examined showed segregation. These included allelic variations for *Yr15* for yellow rust resistance, *Lr34 and Lr21* for leaf rust resistance, *Tsn1* for tan spot sensitivity and *Cre8* for nematode resistance. Two allelic variations for *Yr15* were examined including the causal *Yr15_Kin2* (Klymiuk et al. 2019) and the linked *Yr15_R8* (Ramirez-Gonzalez et al. 2015) polymorphisms. Although showing slightly different distributions, both *Yr15_Kin2* and *Yr15_R8* showed low frequencies of the favorable resistant *+Yr15* allele across the breeding periods (0 – 23%). The distribution is however different for *Lr34* (TCC InDel; Lagudah et al. 2009) and *Tsn1* (Tsn1+/Null; Faris et al. 2010) which both showed pre-dominant and increasing frequencies of their resistant alleles across the breeding periods. The resistant *+Lr21* gene allele was only present in one breeding line and the resistant *+Cre8* allele frequency was stably low (2 - 9%) across the breeding periods.

#### Flowering time genes

EA wheat germplasm mainly comprises photoperiod insensitive spring wheat genotypes which are likely fixed for vernalization and photoperiod insensitivity gene alleles. We, therefore, focused on the *earliness per se* locus, *Eps-D1*, and its homoeologous locus *Eps-B1*, which also account for variation in flowering time irrespective of vernalization and photoperiod requirements (Zikhali et al. 2016; Alvarez et al. 2016). A large chromosomal deletion (including *TaElf3-D1*) and a SNP variation in *TaElf3-D1* have been linked to the *Eps-D1* effect (Zikhali et al. 2016). The chromosomal deletion is flanked by the *TaBradi2g14790* marker. We observed a near-balanced allele frequency distribution for the chromosomal deletion and wild-type (non-deleted) alleles across the breeding periods (Figure S4, Online Resource 2). This deletion variation was also confirmed by the fact that lines showing the chromosomal deletion showed no amplification for *TaElf3-D1* (Figure S5, Online Resource 2). The *Elf3-B1* SNP variation linked to the *Eps-B1* locus was also segregating in the EA population. We observed higher frequencies of the favorable early-flowering *Elf3-B1* allele compared to the late-flowering alleles across the breeding periods with an increasing trend observed from cultivars released from 1970 to modern breeding lines (Figure S4, Online Resource 2).

### Field effect of allelic variations in plant height and grain size genes

We evaluated the phenotypic effects of the allelic variations in plant height and grain size genes in field trials conducted in Ethiopia and Kenya. From these trials, we confirmed the effect of the Green Revolution *Rht-B1b* and *Rht-D1b* semi-dwarfing alleles in reducing plant height. The semi-dwarf alleles of both genes are estimated to significantly (*P* < 0.0001) reduced height by 13.50 (±1.55) cm and 14.93 (±2.23) cm, respectively (Table 1). The *Rht-B1* 160 bp InDel variation did not show any significant effect on plant height in the EA population. The *Rht-B1* 190 bp InDel was almost fixed in the population (99.15%) as such its effect could not be statistically tested. We similarly examined the effect of seven segregating grain size genes including *TaGS-D1, TaCKX-D1, TaCwi-4A, TaSus1-7B, TaGS5-A1, TEF-7A* and *TaGS1a*. Of these genes, only *TaCKX-D1* showed significant (*P* < 0.0001) effect on TKW with the deletion *TaCKX-D1a* allele estimated to increase TKW by 14.27 (±1.99) g (Table 1). However, *TaCKX-D1* only segregates in the Ethiopian population and is fixed for the favorable higher-TKW allele in the Kenyan population.

**Table 1:**
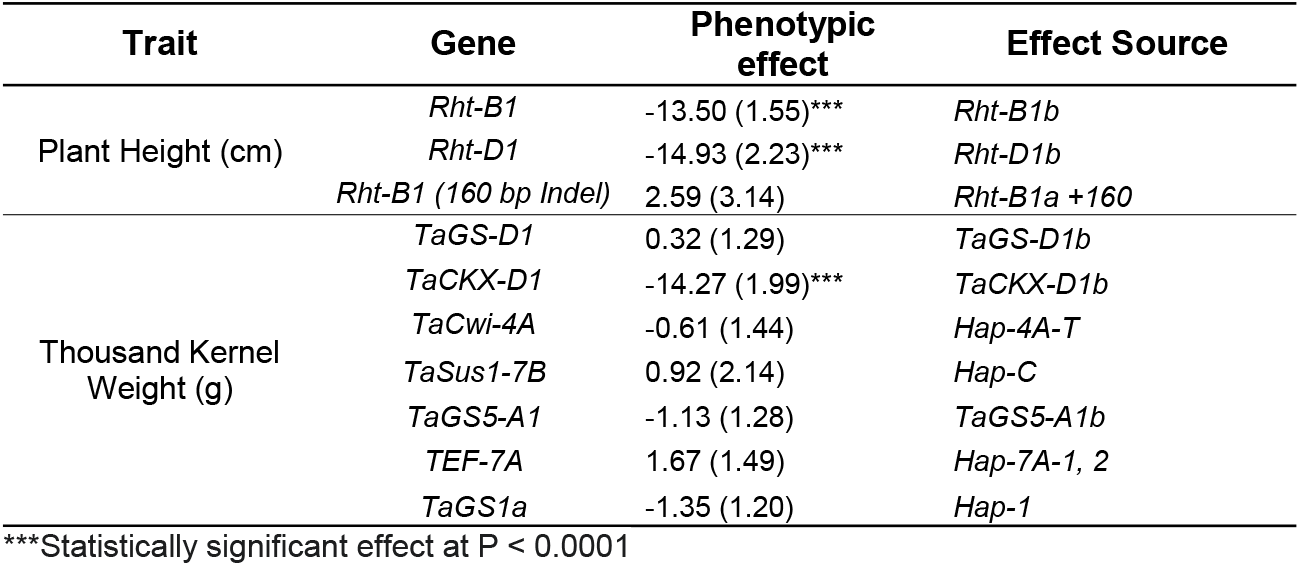
Effects of *Rht-1* and grain size genes on plant height and TKW.

| Trait | Gene | Phenotypic effect | Effect Source |
| --- | --- | --- | --- |
| Plant Height (cm) | *Rht-B1* | -13.50 (1.55)*** | *Rht-B1b* |
|  | *Rht-D1* | -14.93 (2.23)*** | *Rht-D1b* |
|  | *Rht-B1 (160 bp Indel)* | 2.59 (3.14) | *Rht-B1a +160* |
| Thousand Kernel Weight (g) | *TaGS-D1* | 0.32 (1.29) | *TaGS-D1b* |
|  | *TaCKX-D1* | -14.27 (1.99)*** | *TaCKX-D1b* |
|  | *TaCwi-4A* | -0.61 (1.44) | *Hap-4A-T* |
|  | *TaSus1-7B* | 0.92 (2.14) | *Hap-C* |
|  | *TaGS5-A1* | -1.13 (1.28) | *TaGS5-A1b* |
|  | *TEF-7A* | 1.67 (1.49) | *Hap-7A-1, 2* |
|  | *TaGS1a* | -1.35 (1.20) | *Hap-1* |

\*\*\*Statistically significant effect at $P < 0.0001$

## Discussions

We examined the allelic diversity of 42 genes affecting important agronomic, disease resistance, grain size and end-use quality traits in EA wheat population. Most of these genes segregate in the EA population suggesting the presence of favorable and unfavorable alleles, although in different proportions. These highlight opportunities for wheat improvement in EA by using marker-assisted breeding to increase the frequencies of favorable gene alleles.

### Opportunities to introduce major but missing genes in EA germplasm

Some of the major genes missing in EA wheat population include *Sr26, Yr5*, and *Fhb1*. Rusts (stem, yellow and leaf) are major constraints to wheat production in EA. Kenya and Ethiopia are hotspots for highly virulent and rapidly evolving races of the stem rust pathogen (*Puccinia graminis f. sp. tritici*). This includes the Ug99 race (TTKSK) which was first discovered in Uganda in 1998 (Pretorius et al. 2000; Singh et al. 2011) and was virulent to most of the stem rust resistance genes deployed in global wheat germplasm at that time. About four additional variants of Ug99 each with increasing virulence to more stem rust resistance genes and five new non-Ug99 races have since been reported in Kenya and Ethiopia (Singh et al. 2015; Bhavani et al. 2019). *Sr26*, a gene introgressed from *Agropyron elongatum* (Liu et al. 2010), is one of very few resistance genes still effective against all known Ug99 variants. However, its use in global germplasm has been limited as a result of an associated yield penalty caused by linkage drag (Liu et al. 2010). This yield penalty possibly accounts for the lack of *Sr26* in EA wheat population, although there is no evidence to suggest that *Sr26* was ever introduced or tested under EA conditions. Lines with smaller introgression of *Sr26* without the yield loss penalty (Dundas et al. 2007) have been developed, thus providing an opportunity for deploying and evaluating the effect of *Sr26* on rust resistance and yield under EA conditions.

Similarly, *Yr5* is one of very few genes conferring resistance against a wide range of virulent strains of the yellow rust pathogen (*Puccinia striiformis f. sp.tritici*; Marchal et al. 2018). Unlike *Sr26, Yr5* is currently used in many breeding programs including Europe and has no associated yield penalty. There should, therefore, be no disincentive against the use of *Yr5* in EA germplasm. The infrequent occurrence of Fusarium Head Blight (FHB) in EA might account for the lack of *Fhb1* in EA germplasm. Nonetheless, FHB still causes significant yield loss and poses health risks due to the production of hazardous mycotoxins. There is, therefore, an opportunity to anticipatorily breed for resistance against the sporadic occurrence of FHB.

It is worth noting that we only examined disease resistance genes for which KASP markers have been developed. We cannot, therefore, exclude the presence of other functional resistance genes, particularly for stem and yellow rust, in the EA wheat population. The development and public deposition of more breeder friendly KASP marker assays for resistance genes will facilitate the examination and deployment of these genes in global germplasm. Also, the use of KASP markers to rapidly verify the absence of known resistance genes in otherwise resistant germplasm will highlight the presence and facilitate the identification of novel resistance genes in such resistant germplasm.

### Allele frequency changes across breeding periods and balancing selection for optimizing traits expression

We also examined changes in allele frequency distribution across different breeding periods for segregating genes alleles. This analysis highlights the progress made in increasing the favorable alleles of some genes (including *Rht-1, Glu-D1, Pin-A, Tsn1* and *Lr34*) across the breeding periods. For genes like *Rht-B1* and *Lr34* whose effects are easily observed, the progressive increase in favorable allele frequencies might be due to direct phenotypic selection. The advent of breeder-friendly molecular markers in wheat breeding might have also accelerated the selection of these alleles. For instance, the drastic increase in the semi-dwarf (favorable) allele of *Rht-B1* in the lines released between 2001 – 2017 coincided with the development of “perfect” molecular markers for the *Rht-1* genes by Ellis et al (2002).

Our study also highlights the progressive loss of the favorable alleles of some genes: *Rht-D1, Pinb-D1, TaMKK-3A*. While it is important to work towards increasing the favorable alleles of these genes, it is important to bear in mind that the overall goal of an effective breeding program should be on traits improvement (i.e. improving favorable trait expression) rather than “allele” improvement *per se* (i.e. increasing favorable gene allele frequency). This is particularly important for multigenic traits where interactions (additive or epistatic) between the different genes controlling such traits are more important in determining the trait expression than the individual gene actions. Balancing the selection of interacting gene alleles is therefore important for fine-tuning and optimizing the expression of such traits for different environments and end-uses. We found evidence for such balancing selection in EA germplasm. For instance, *Rht-B1* and *Rht-D1* additively control plant height, with plants having semi-dwarf alleles of both genes showing severely reduced plant height (dwarf). This dwarf phenotype is, however, not considered beneficial in Kenya and Ethiopia because it reduces the amount of straws farmers can harvest for feeding livestock. Consequently, we observed that increase in the *Rht-B1b* allele is accompanied by a decrease in *Rht-D1b* allele. This is also true for *TaMKK3-A* and *TaMFT-A1* which both control seed dormancy levels in an additive manner. Although the dormancy-promoting allele frequencies of both genes showed an overall decreasing trend, we observed that high frequency of the dormancy-promoting *TaMFT1* allele is balanced by low frequency of the dormancy-promoting *TaMKK3-A* allele. Only 13% of lines examined contains dormancy-promoting alleles of both genes. This balanced selection ensures that an optimum level of dormancy is expressed that protects against PHS without impairing uniform seed germination.

### Importance of validating population-specific effect of beneficial genes

Some of the markers investigated in this study, particularly for grain size genes, were identified by association analyses in wheat populations that are genetically distinct to the EA wheat population used in this study. It was therefore important to validate the effect of such genes in EA germplasm under local field conditions. Out of the seven grain-size related genes segregating in the EA wheat population, only *TaCKX-D1* showed a significant effect on TKW. It is important to note that the polymorphism reported in these genes were only shown to be associated with, but were not validated as being causal of the natural variation in TKW (Zhang et al. 2012; Guo et al. 2013; Hou et al. 2014; Zheng et al. 2014; Jiang et al. 2015; Wang et al. 2016). It is, therefore, possible that the lack of phenotypic effect in our study might be due to difference in the haplotype structure between the EA wheat population and the original study population used to identify these markers. Such haplotype differences would not have been picked up through single-marker analysis for each gene as was done our study. Grain-size traits are also highly environmental dependent (Brinton and Uauy 2019), and it is possible that differences in environmental conditions might also have contributed to the lack of effects of some of the grain size genes examined. Similarly, Sukumaran et al (2018) did not find consistent significant effect of a reported functional allelic variation of an important grain size gene, *TaGW2-A*, on TKW when tested under 10 environments. On the other hand, causal polymorphisms in *Rht-B1* and *Rht-D1* which have been widely validated across global germplasm showed significant effects in our study. We therefore propose that haplotype-based analysis (involving the use of markers defining haplotype block around a gene of interest) should be used in validating the effect of associated genes under local growth conditions before using such genes for marker-assisted breeding.

### Outlook

Knowledge of the allelic diversity currently used or missing in EA wheat population is important in informing strategies for wheat breeding in Kenya and Ethiopia. This is particularly important for combating diseases that are critically limiting wheat production in both countries. To this end, we are currently working towards introducing beneficial alleles of *Yr5* (missing), *Yr15* (low frequency), *Sr26* (missing) and *Sr22* in combination in a common locally adapted EA wheat line. We will also be introducing functional *Fhb1* (missing) alleles to this background to future proof the developed lines against the sporadic occurrence of FHB.

## Supporting information

Online Resource 1

Online Resource 2

## Declarations

### Contributions

OS, CU, NY and RW designed the experiments. MW, ZT, LM, HZ, MR and OS performed the experiments (sampling, genotyping, and field work) and analyzed the results. MW, ZT and OS wrote the manuscript. All authors read and approved the manuscript.

