## Supplementary material for "Allelic diversity study of functional genes in East Africa bread wheat highlights opportunities for genetic improvement": Online Resource 2

### Slide 1
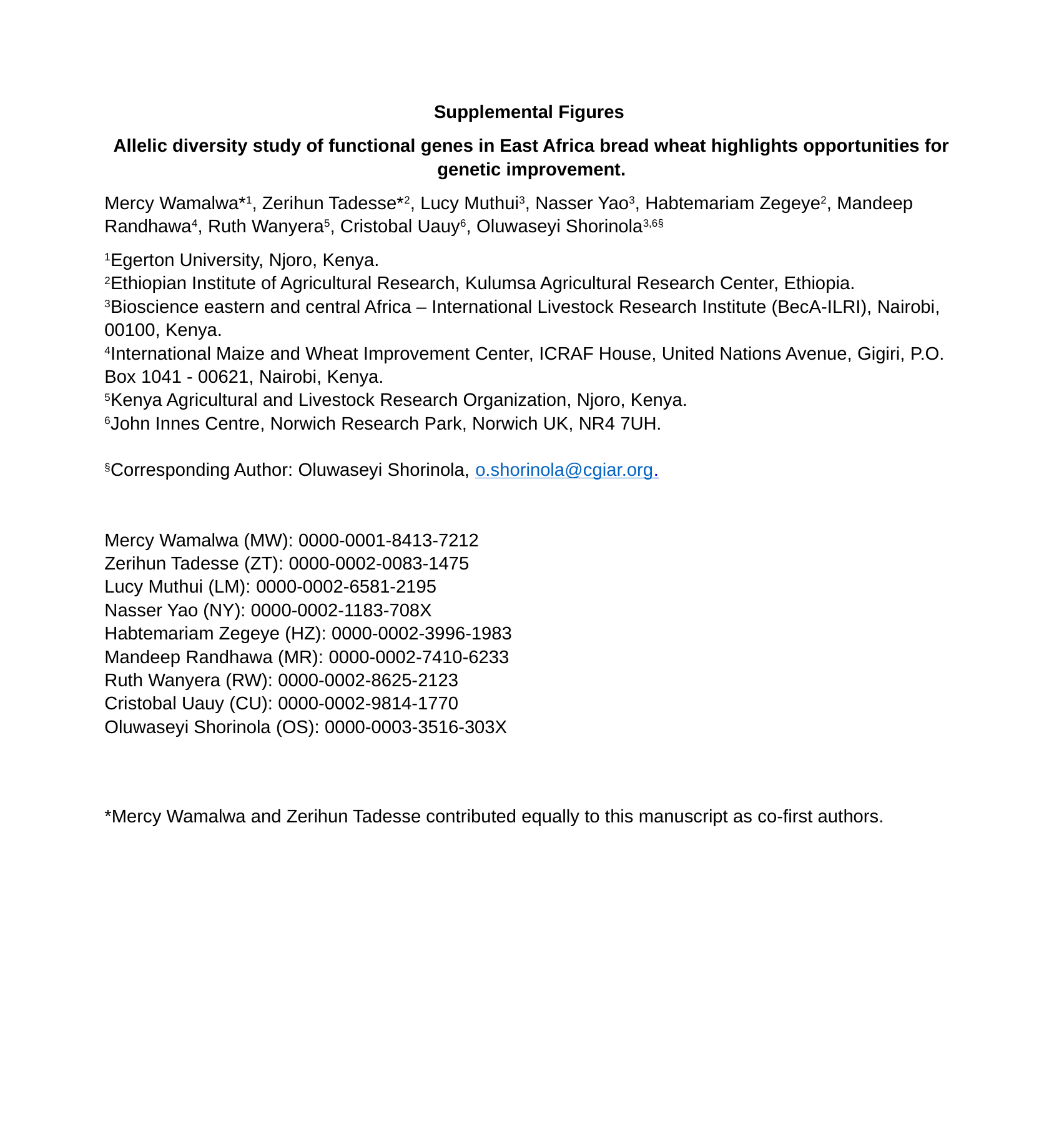

Supplemental Figures
Allelic diversity study of functional genes in East Africa bread wheat highlights opportunities for genetic improvement.
Mercy Wamalwa*^1^, Zerihun Tadesse*^2^, Lucy Muthui^3^, Nasser Yao^3^, Habtemariam Zegeye^2^, Mandeep Randhawa^4^, Ruth Wanyera^5^, Cristobal Uauy^6^, Oluwaseyi Shorinola^3,6§^
^1^Egerton University, Njoro, Kenya.
^2^Ethiopian Institute of Agricultural Research, Kulumsa Agricultural Research Center, Ethiopia.
^3^Bioscience eastern and central Africa – International Livestock Research Institute (BecA-ILRI), Nairobi, 00100, Kenya.
^4^International Maize and Wheat Improvement Center, ICRAF House, United Nations Avenue, Gigiri, P.O. Box 1041 - 00621, Nairobi, Kenya.
^5^Kenya Agricultural and Livestock Research Organization, Njoro, Kenya.
^6^John Innes Centre, Norwich Research Park, Norwich UK, NR4 7UH.
Mercy Wamalwa (MW): 0000-0001-8413-7212
Zerihun Tadesse (ZT): 0000-0002-0083-1475
Lucy Muthui (LM): 0000-0002-6581-2195
Nasser Yao (NY): 0000-0002-1183-708X
Habtemariam Zegeye (HZ): 0000-0002-3996-1983
Mandeep Randhawa (MR): 0000-0002-7410-6233
Ruth Wanyera (RW): 0000-0002-8625-2123
Cristobal Uauy (CU): 0000-0002-9814-1770
Oluwaseyi Shorinola (OS): 0000-0003-3516-303X
*Mercy Wamalwa and Zerihun Tadesse contributed equally to this manuscript as co-first authors.

### Slide 2
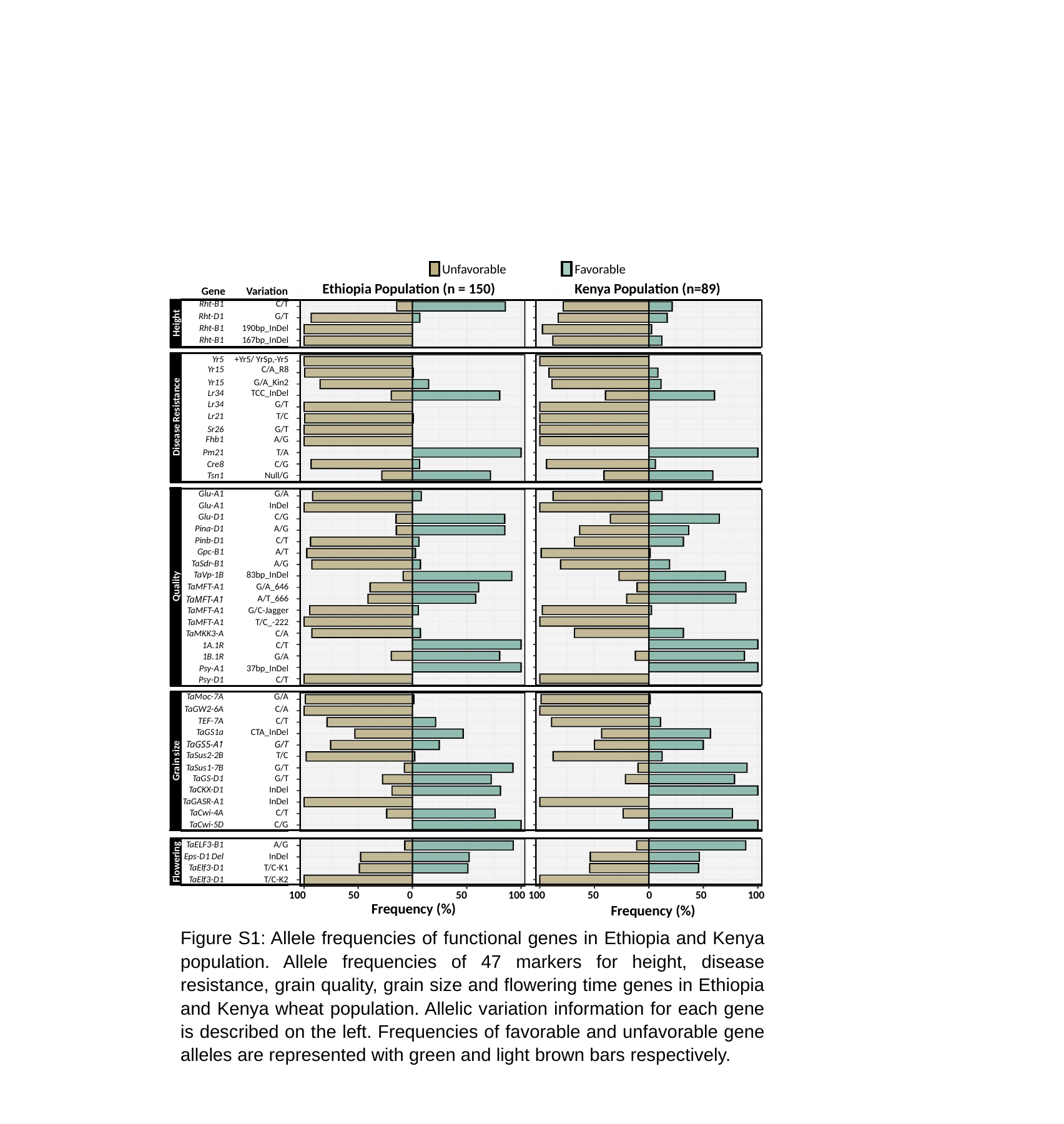

Unfavorable
Favorable
Ethiopia Population (n = 150)
Kenya Population (n=89)
Gene
Variation
| Rht-B1 | C/T |
| --- | --- |
| Rht-D1 | G/T |
| Rht-B1 | 190bp\_InDel |
| Rht-B1 | 167bp\_InDel |
Height
| Yr5 | +Yr5/ YrSp,-Yr5 |
| --- | --- |
| Yr15 | C/A\_R8 |
| Yr15 | G/A\_Kin2 |
| Lr34 | TCC\_InDel |
| Lr34 | G/T |
| Lr21 | T/C |
| Sr26 | G/T |
| Fhb1 | A/G |
| Pm21 | T/A |
| Cre8 | C/G |
| Tsn1 | Null/G |
Disease Resistance
| Glu-A1 | G/A |
| --- | --- |
| Glu-A1 | InDel |
| Glu-D1 | C/G |
| Pina-D1 | A/G |
| Pinb-D1 | C/T |
| Gpc-B1 | A/T |
| TaSdr-B1 | A/G |
| TaVp-1B | 83bp\_InDel |
| TaMFT-A1 | G/A\_646 |
| TaMFT-A1 | A/T\_666 |
| TaMFT-A1 | G/C-Jagger |
| TaMFT-A1 | T/C\_-222 |
| TaMKK3-A | C/A |
| 1A.1R | C/T |
| 1B.1R | G/A |
| Psy-A1 | 37bp\_InDel |
| Psy-D1 | C/T |
Quality
| TaMoc-7A | G/A |
| --- | --- |
| TaGW2-6A | C/A |
| TEF-7A | C/T |
| TaGS1a | CTA\_InDel |
| TaGS5-A1 | G/T |
| TaSus2-2B | T/C |
| TaSus1-7B | G/T |
| TaGS-D1 | G/T |
| TaCKX-D1 | InDel |
| TaGASR-A1 | InDel |
| TaCwi-4A | C/T |
| TaCwi-5D | C/G |
Grain size
| TaELF3-B1 | A/G |
| --- | --- |
| Eps-D1 Del | InDel |
| TaElf3-D1 | T/C-K1 |
| TaElf3-D1 | T/C-K2 |
Flowering
100
50
0
50
100
100
50
0
50
100
Frequency (%)
Frequency (%)
Figure S1: Allele frequencies of functional genes in Ethiopia and Kenya population. Allele frequencies of 47 markers for height, disease resistance, grain quality, grain size and flowering time genes in Ethiopia and Kenya wheat population. Allelic variation information for each gene is described on the left. Frequencies of favorable and unfavorable gene alleles are represented with green and light brown bars respectively.

### Slide 3
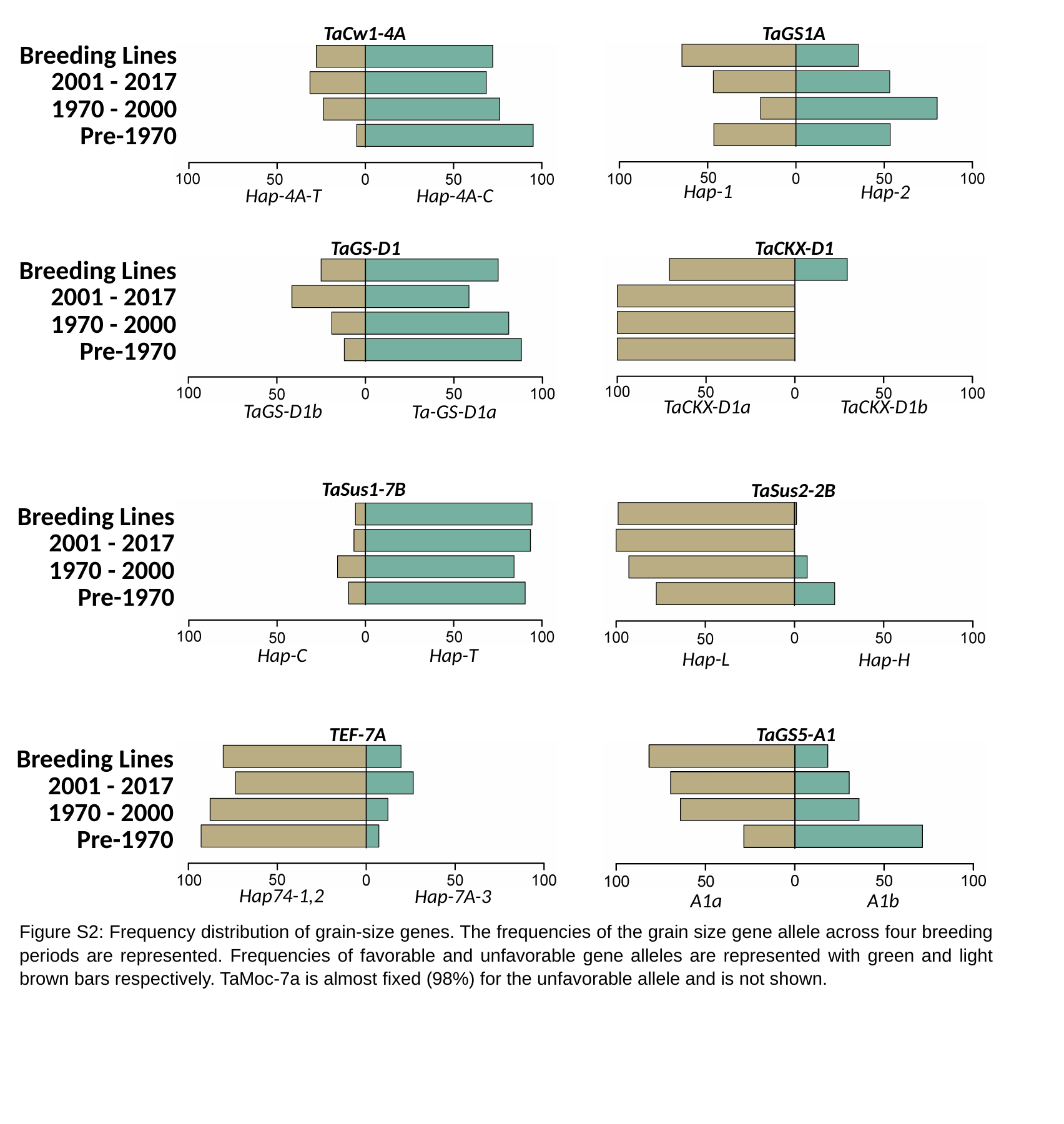

TaCw1-4A
TaGS1A
Breeding Lines
2001 - 2017
1970 - 2000
Pre-1970
Hap-1
Hap-2
Hap-4A-T
Hap-4A-C
TaGS-D1
TaCKX-D1
Breeding Lines
2001 - 2017
1970 - 2000
Pre-1970
TaCKX-D1a
TaCKX-D1b
TaGS-D1b
Ta-GS-D1a
TaSus1-7B
TaSus2-2B
Breeding Lines
2001 - 2017
1970 - 2000
Pre-1970
Hap-C
Hap-T
Hap-L
Hap-H
TEF-7A
TaGS5-A1
Breeding Lines
2001 - 2017
1970 - 2000
Pre-1970
Hap74-1,2
Hap-7A-3
A1a
A1b
Figure S2: Frequency distribution of grain-size genes. The frequencies of the grain size gene allele across four breeding periods are represented. Frequencies of favorable and unfavorable gene alleles are represented with green and light brown bars respectively. TaMoc-7a is almost fixed (98%) for the unfavorable allele and is not shown.

### Slide 4
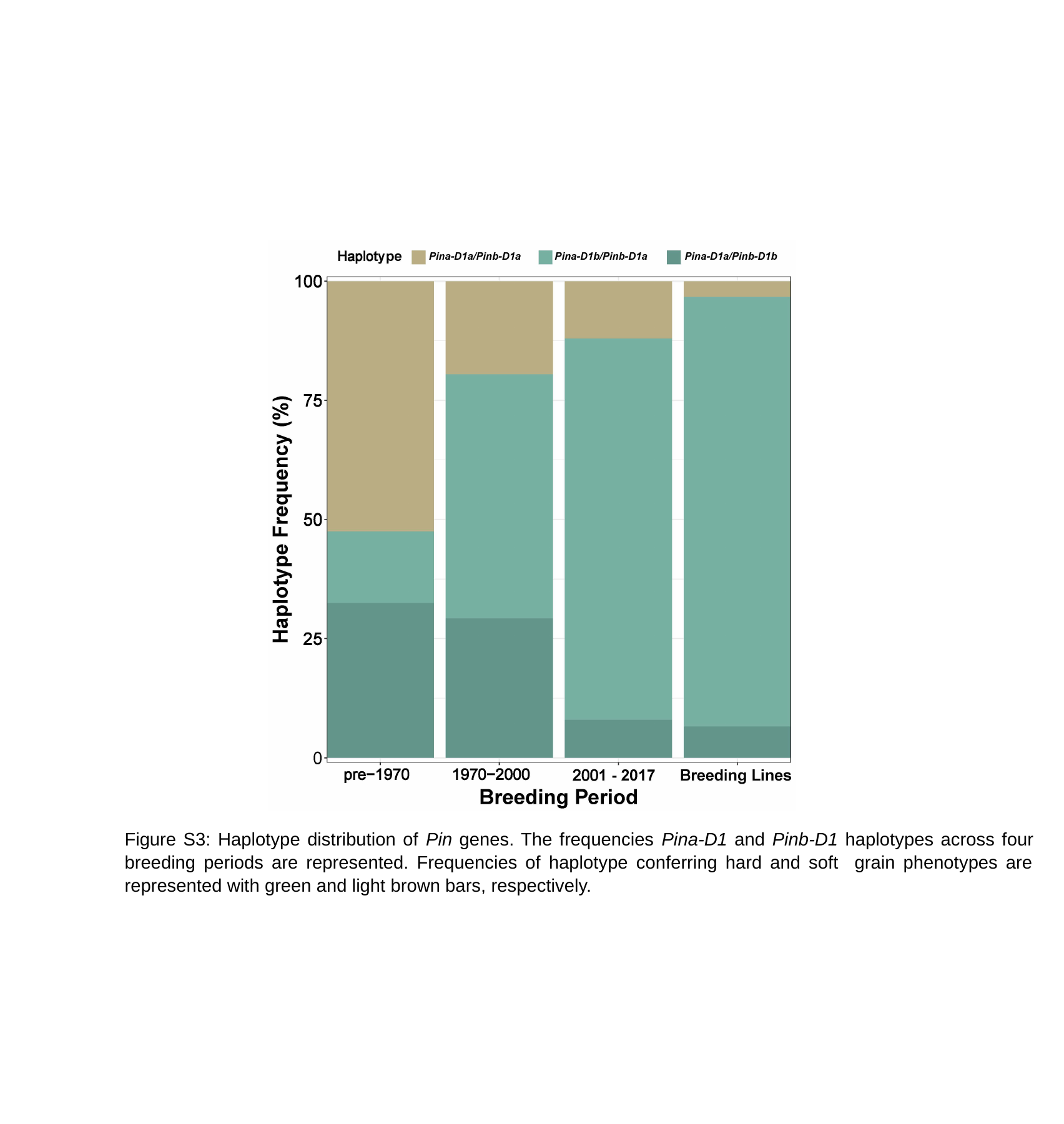

Figure S3: Haplotype distribution of Pin genes. The frequencies Pina-D1 and Pinb-D1 haplotypes across four breeding periods are represented. Frequencies of haplotype conferring hard and soft grain phenotypes are represented with green and light brown bars, respectively.

### Slide 5
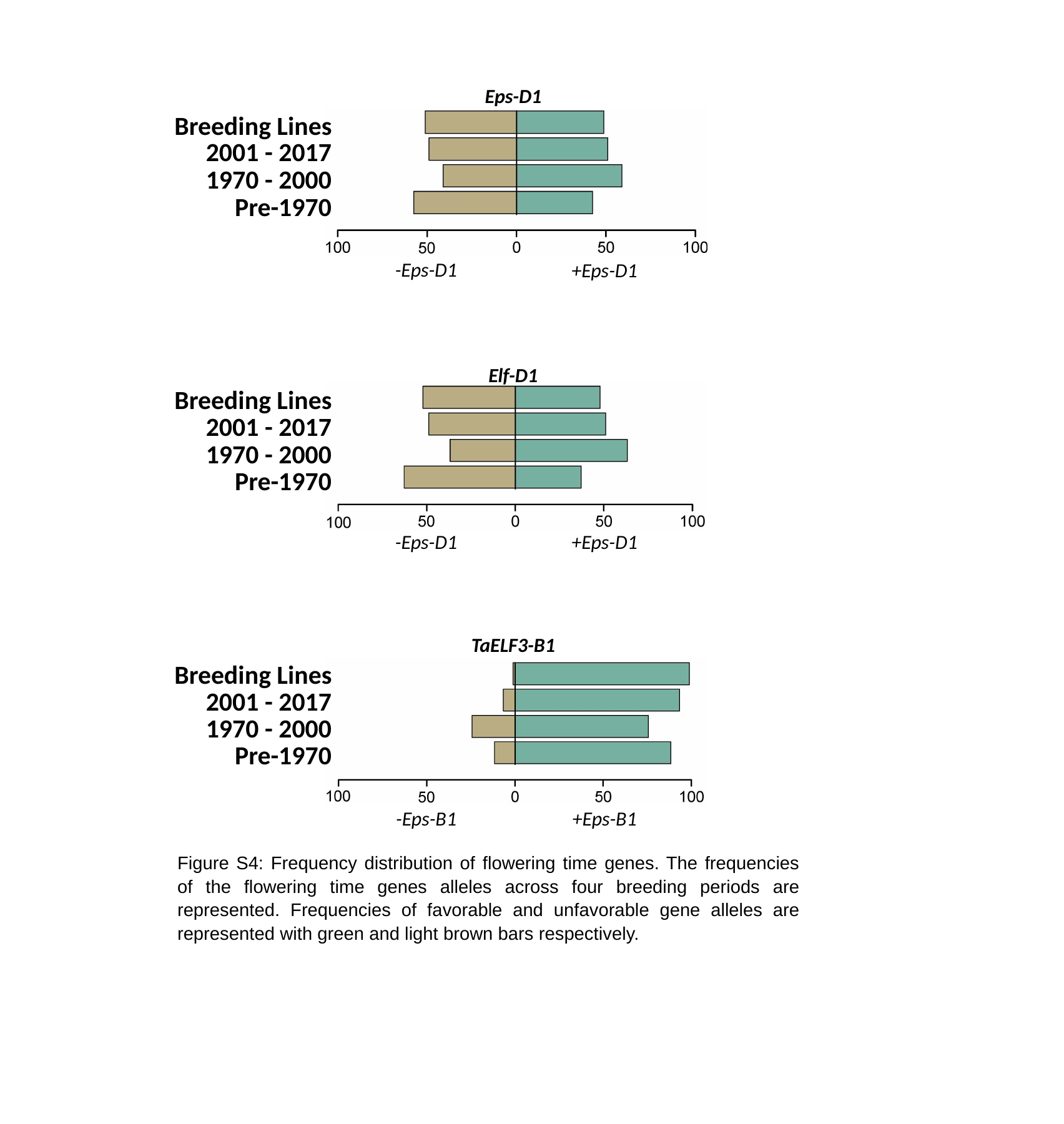

Eps-D1
Breeding Lines
2001 - 2017
1970 - 2000
Pre-1970
-Eps-D1
+Eps-D1
Elf-D1
Breeding Lines
2001 - 2017
1970 - 2000
Pre-1970
-Eps-D1
+Eps-D1
TaELF3-B1
Breeding Lines
2001 - 2017
1970 - 2000
Pre-1970
-Eps-B1
+Eps-B1
Figure S4: Frequency distribution of flowering time genes. The frequencies of the flowering time genes alleles across four breeding periods are represented. Frequencies of favorable and unfavorable gene alleles are represented with green and light brown bars respectively.

### Slide 6
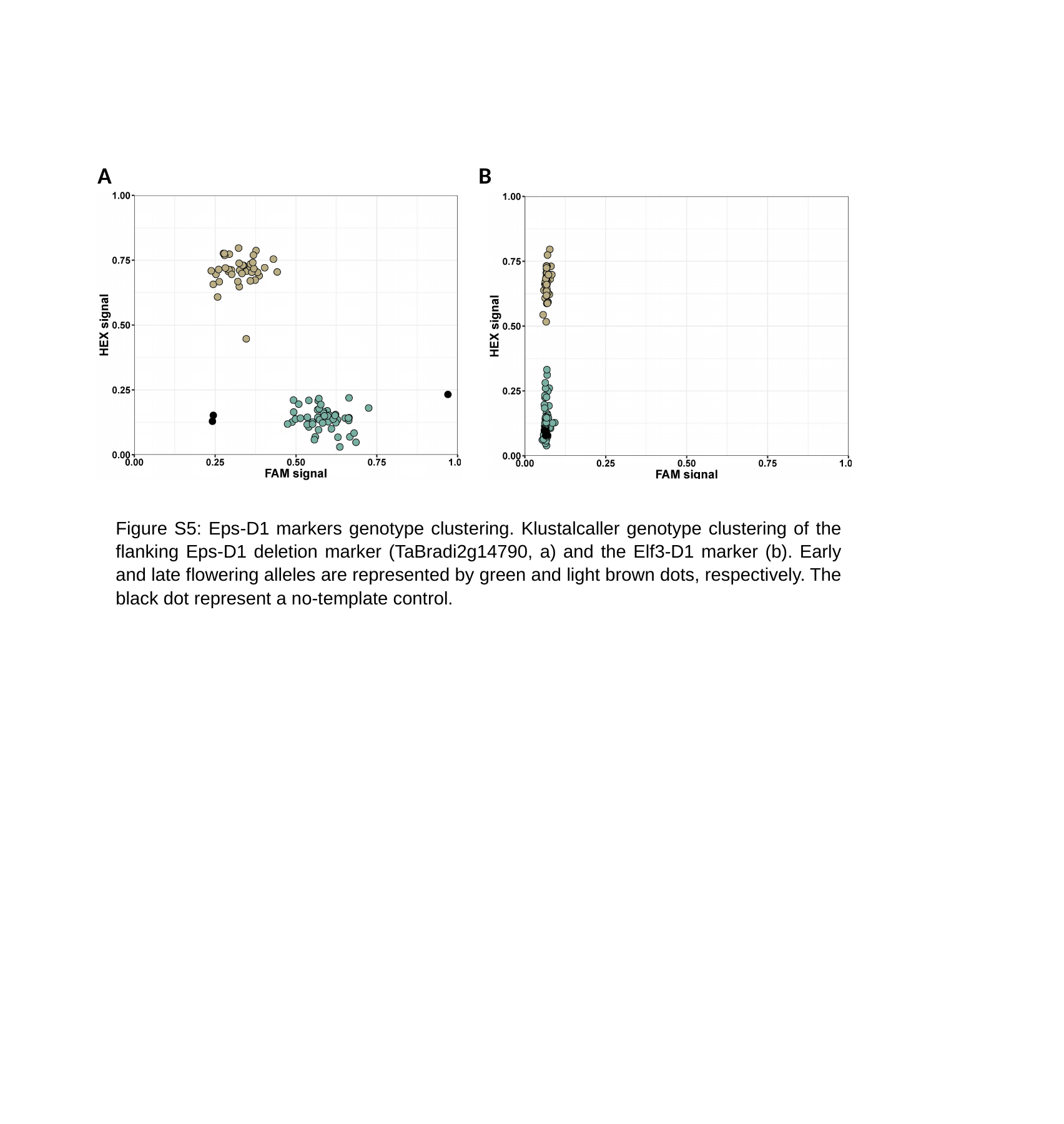

A
B
Figure S5: Eps-D1 markers genotype clustering. Klustalcaller genotype clustering of the flanking Eps-D1 deletion marker (TaBradi2g14790, a) and the Elf3-D1 marker (b). Early and late flowering alleles are represented by green and light brown dots, respectively. The black dot represent a no-template control.
